# The cellular NMD pathway restricts Zika virus infection and is targeted by the viral capsid protein

**DOI:** 10.1101/290296

**Authors:** Krystal A. Fontaine, Kristoffer E. Leon, Mir M. Khalid, Sakshi Tomar, David Jimenez-Morales, Mariah Dunlap, Julia A. Kaye, Priya S. Shah, Steve Finkbeiner, Nevan J. Krogan, Melanie Ott

## Abstract

Zika virus (ZIKV) infection of neural progenitor cells (NPCs) *in utero* is associated with neurological disorders, such as microcephaly, but a detailed molecular understanding of ZIKV-induced pathogenesis is lacking. Here we show that *in vitro* ZIKV infection of human cells, including NPCs, causes disruption of the nonsense-mediated mRNA decay (NMD) pathway. NMD is a cellular mRNA surveillance mechanism that is required for normal brain size in mice. Using affinity purification-mass spectrometry, we identified multiple cellular NMD factors that bind to the viral capsid protein, including the central NMD regulator up-frameshift protein 1 (UPF1). Endogenous UPF1 interacted with the ZIKV capsid protein in co-immunoprecipitation experiments, and capsid expression post-transcriptionally downregulated UPF1 protein levels, a process that we confirmed occurs during ZIKV infection. Cellular fractionation studies show that the ZIKV capsid protein specifically targets nuclear UPF1 for degradation via the proteasome. A further decrease in UPF1 levels by RNAi significantly enhanced ZIKV infection in NPC cultures, consistent with a model in which NMD restricts ZIKV infection in the fetal brain. We propose that ZIKV, via the capsid protein, has evolved a strategy to lower UPF1 levels and dampen antiviral activities of NMD, which in turn contributes to neuropathology *in vivo*.

**Importance:** Zika virus (ZIKV) is a significant global health threat, as infection has been linked to serious neurological complications, including microcephaly. Using a human stem cell-derived neural progenitor model system, we find that a critical cellular quality control process called the nonsense-mediated mRNA decay (NMD) pathway is disrupted during ZIKV infection. Importantly, disruption of the NMD pathway is a known cause of microcephaly and other neurological disorders. We further identify an interaction between the capsid protein of ZIKV and up-frameshift protein 1 (UPF1), the master regulator of NMD, and show that ZIKV capsid targets UPF1 for degradation. Together, these results offer a new mechanism for how ZIKV infection can cause neuropathology in the developing brain.

## Introduction

ZIKV is a mosquito-borne RNA virus that belongs to the *Flaviviridae* family. First isolated in Uganda in 1947, ZIKV remained relatively obscure for decades following its discovery because infection was associated with only mild disease. However, more severe clinical manifestations, including microcephaly, have been observed during the recent spread of ZIKV through the Americas (1). ZIKV infection induces cell cycle arrest and apoptosis in neural progenitor cells (NPCs) in *in vitro* studies and *in vivo* mouse models, with the latter resulting in cortical thinning and microcephaly (2–6). While it is now established that ZIKV infection during pregnancy is a causative agent of microcephaly (7), the molecular mechanisms underlying ZIKV-induced neuropathogenesis remain largely unknown.

Similar to other flaviviruses, ZIKV contains a single-stranded, positive-sense RNA genome of ∼11 kb in size. The genome encodes a single polyprotein that is post-translationally processed by both host and viral proteases to produce 3 structural and 7 nonstructural proteins (8, 9). The flavivirus capsid, which is the first protein encoded in the genome, is a major structural element required for the encapsidation of the RNA genome during virion assembly (10). While flavivirus replication is known to occur in the cytoplasm, a significant portion of the viral capsid protein localizes to the nucleus during infection (10, 11). Although the role of nuclear capsid during infection is less clear, several functions have been suggested. The capsid protein from dengue virus, a close relative of ZIKV, binds to core histones and inhibits nucleosome formation, thus implicating the protein in altering host gene expression (12). Furthermore, several flavivirus capsid proteins, including ZIKV capsid, localize to the nucleolus, with many interacting with nucleolar proteins to promote viral particle production (13–16).

The nonsense-mediated mRNA decay (NMD) pathway was initially discovered as a highly conserved quality control system that destroys transcripts containing premature termination codons (PTCs)(17). Following splicing of pre-mRNAs, a multi-subunit protein complex called the exon-junction complex (EJC) is deposited onto mRNAs near the sites of exon-exon junctions. If a PTC is found ∼50–55 nucleotides upstream of an EJC, the mRNA will be subjected to NMD-mediated degradation initiated by the recruitment of the RNA helicase up-frameshift protein 1 (UPF1). UPF1 plays a central role in the NMD pathway by linking the translation termination event to the assembly of a surveillance complex, resulting in NMD activation (18). Interestingly, microcephaly has been associated with genetic mutations that result in the impairment of the NMD pathway. While knockout of Upf1 and other NMD factors is embryonic lethal in mice (19), mice haploinsufficient for the EJC components Magoh, Rbm8a, and Eif4a3 exhibit aberrant neurogenesis and microcephaly (20–22).

In addition to PTC-containing transcripts, it is now known that the NMD pathway recognizes a broader range of RNA substrates. Notably, the NMD controls the “normal” expression of ∼10% of the cellular transcriptome and is regarded as a post-transcriptional mechanism of gene regulation (23). Furthermore, the NMD pathway also regulates viral infections. While it was first reported that UPF1 promotes the infectivity of HIV-1 progeny virions (24), replication of several human RNA viruses, including human T-cell lymphotropic virus type 1 (HTLV-1), Semliki Forest virus and Sindbis virus, is enhanced following UPF1 knockdown, implicating UPF1 and the NMD pathway, either directly or indirectly, in the host antiviral response (25–28). As ZIKV infection and NMD impairment both promote microcephaly development, and we previously described disruption of the NMD pathway in cells infected with a related flavivirus, the hepatitis C virus (HCV) (29), we hypothesized that ZIKV infection manipulates the cellular NMD pathway, a process contributing to ZIKV-induced neuropathology.

## Results

### The NMD pathway is impaired during ZIKV infection

To determine if ZIKV infection affects NMD, we infected human hepatoma cells (Huh7) and human induced pluripotent stem cell (iPSC)-derived NPCs with ZIKV for 48 h. We isolated total RNA from infected cells and measured mRNA levels of three canonical NMD substrates: asparagine synthetase (ASNS), cysteinyl-tRNA synthetase (CARS), and SR protein SC35 (29). ASNS, CARS, and SC35 transcripts were significantly elevated in Huh7 cells and NPCs following infection with Asian lineage ZIKV strain P6-740 (Fig. 1a). Levels of NMD substrates were also elevated in Huh7 cells infected with the contemporary ZIKV clinical isolate PRVABC59 (Puerto Rico, 2015) (Fig. 1a). We found that the ZIKV-induced increase in NMD transcripts did not reflect a global increase in transcription, as mRNA levels of housekeeping genes, including glyceraldehyde 3-phosphate dehydrogenase (GAPDH), were not altered in infected cells (Fig. 1a). Together, these results indicate that the NMD pathway is impaired in ZIKV-infected cells.

**Figure 1.**
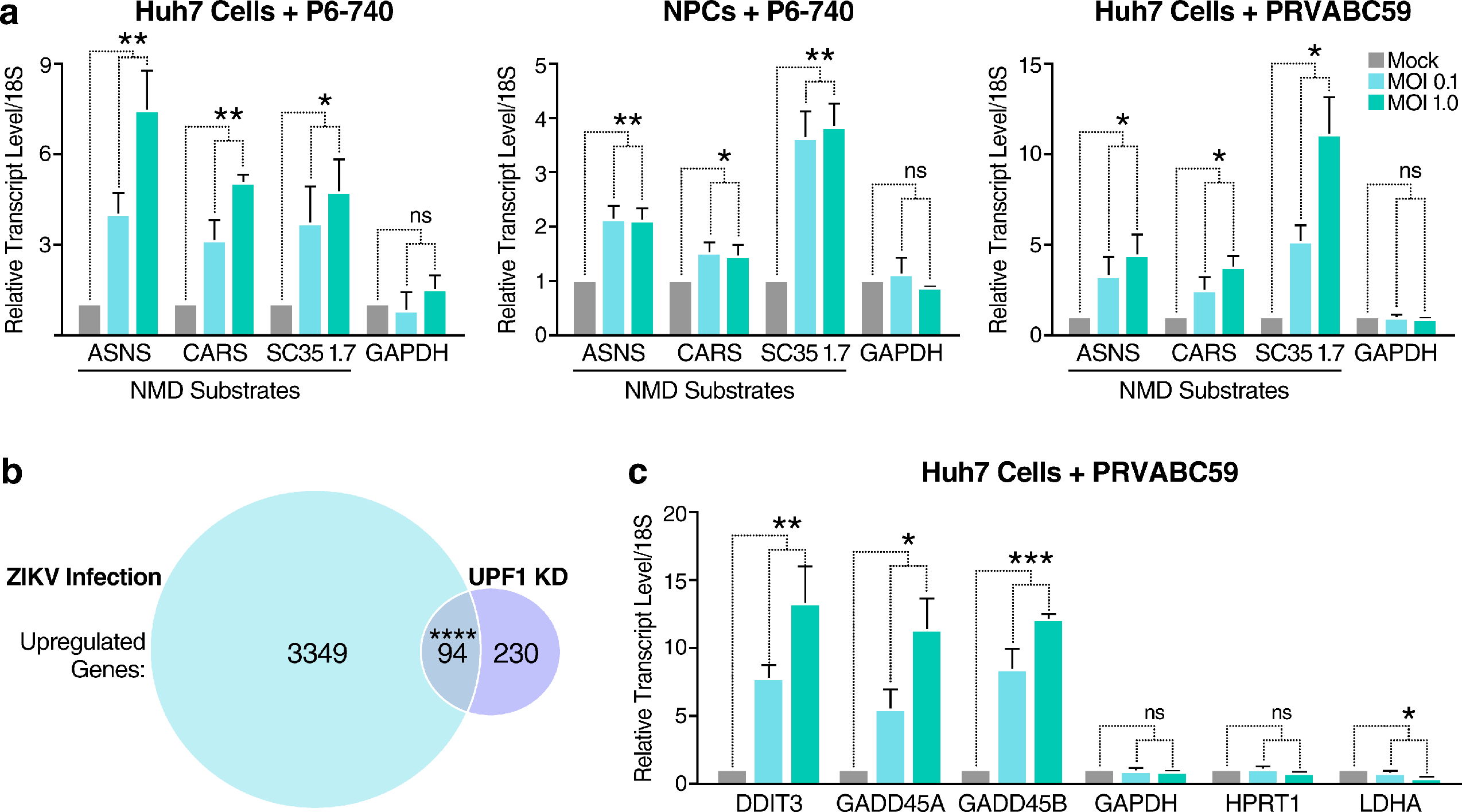
ZIKV infection disrupts the NMD pathway. (a) Transcript levels of NMD substrates and housekeeping genes from Huh7 cells or NPCs mock-infected or infected with ZIKV strain P6-740 or the contemporary clinical isolate PRVABC59. Cells were infected at a multiplicity of infection (MOI) of 0.1 or 1 and harvested at 48 hpi. Data are represented as mean ± s.e.m. *P* values were calculated by unpaired Student’s *t*-test. \**P* ≤ 0.05; \*\**P* ≤ 0.01; ns, not significant. n= 3 independent experiments. (b) Venn diagram showing overlap of significantly upregulated genes associated with ZIKV infection of NPCs and UPF1 knockdown in HeLa cells. RNA-Seq analyses of mock-infected or ZIKV-infected NPCs harvested at 56 hpi and control siRNA-treated or UPF1 siRNA-treated HeLa TO cells harvested at 72 h post-transfection (hpt). The GeneProf hypergeometric probability calculator was then used to generate a hypergeometric *P* value. \*\*\*\**P* ≤ 0.0001. (c) Transcript levels of housekeeping genes and select genes involved in cell cycle growth arrest and apoptosis that were identified in (b). Huh7 cells were mock-infected or infected with ZIKV PRVABC59 at an MOI of 0.1 or 1 and harvested at 48 hpi. Data are represented as mean ± s.e.m. *P* values were calculated by unpaired Student’s *t*-test. \**P* ≤ 0.05; \*\**P* ≤ 0.01; \*\*\**P* ≤ 0.001; ns, not significant. n= 3 independent experiments.

NMD substrates are regulated through the ATP-dependent RNA helicase activity of UPF1, the central regulator of NMD (18). To determine if ZIKV infection broadly affects NMD, we utilized two publicly available RNA sequencing (RNA-Seq) datasets to compare genome-wide transcriptional alterations found during ZIKV infection (6) to those found following UPF1 knockdown (30). As shown in Figure 1b, there is a significant overlap in upregulated genes between these two datasets. Interestingly, several of the overlapping genes are canonical NMD substrates (31–35) involved in cell cycle arrest and induction of apoptosis, two conditions linked to ZIKV-associated neuropathology (5). These genes include DNA damage-inducible transcript 3 (DDIT3) (36) and growth arrest and DNA damage-inducible protein 45 alpha and beta (GADD45A and GADD45B, respectively) (37). Via quantitative real-time RT-PCR, we confirmed that transcripts of each were upregulated following infection of Huh7 cells with ZIKV PRVABC59, while the mRNA levels of the housekeeping genes GAPDH, hypoxanthine phosphoribosyltransferase 1 (HPRT1), and lactate dehydrogenase A (LDHA) were not elevated (Fig. 1c). Combined, these data show that ZIKV infection is associated with dysregulated expression of NMD substrates relevant to ZIKV-mediated neuropathogenesis.

### ZIKV capsid interacts with the NMD pathway

We previously showed that the core protein of HCV, as well as the capsid protein of the related flaviviruses dengue virus and West Nile virus, interact with the host protein within bgcn homolog (WIBG/PYM1), an EJC disassembly factor associated with NMD (29, 38). To examine potential interactions between ZIKV and the NMD pathway, we separately analyzed data generated from an affinity purification-mass spectrometry (AP-MS) screen to specifically query whether the capsid protein of ZIKV interacts with NMD-associated host factors (Shah et al., submitted). ZIKV-host protein-protein interaction (PPI) maps were generated in HEK293T cells using ZIKV proteins from the Ugandan 1947 strain MR 766 or the French Polynesian 2013 strain H/PF/2013 as bait proteins.

From this analysis, we found that ZIKV capsid proteins interacted with several components of the NMD pathway, including multiple members of the EJC complex, as well as UPF1 and UPF3B, the latter an NMD effector that stimulates UPF1 helicase activity (Fig. 2a) (39). Importantly, the NMD host factors that interacted with each of the two different capsid proteins greatly overlapped, revealing that the interaction between capsid and the NMD pathway is conserved across the Asian and African lineages of ZIKV (Fig. 2a). Next, we validated the binding of ZIKV capsid to select NMD host factors by co-immunoprecipitating FLAG-tagged capsid protein with endogenous UPF3B or UPF1 in HEK293T cells. Both UPF3B and UPF1 proteins co-immunoprecipitated with ZIKV capsid, thus confirming the AP-MS results (Fig. 2b,c, respectively). UPF1 interacted with the viral capsid protein independently from its RNA-binding and ATPase/helicase capacities, as ZIKV capsid co-immunoprecipitated with UPF1 mutants deficient in these functions following overexpression (Fig 2d).

**Figure 2.**
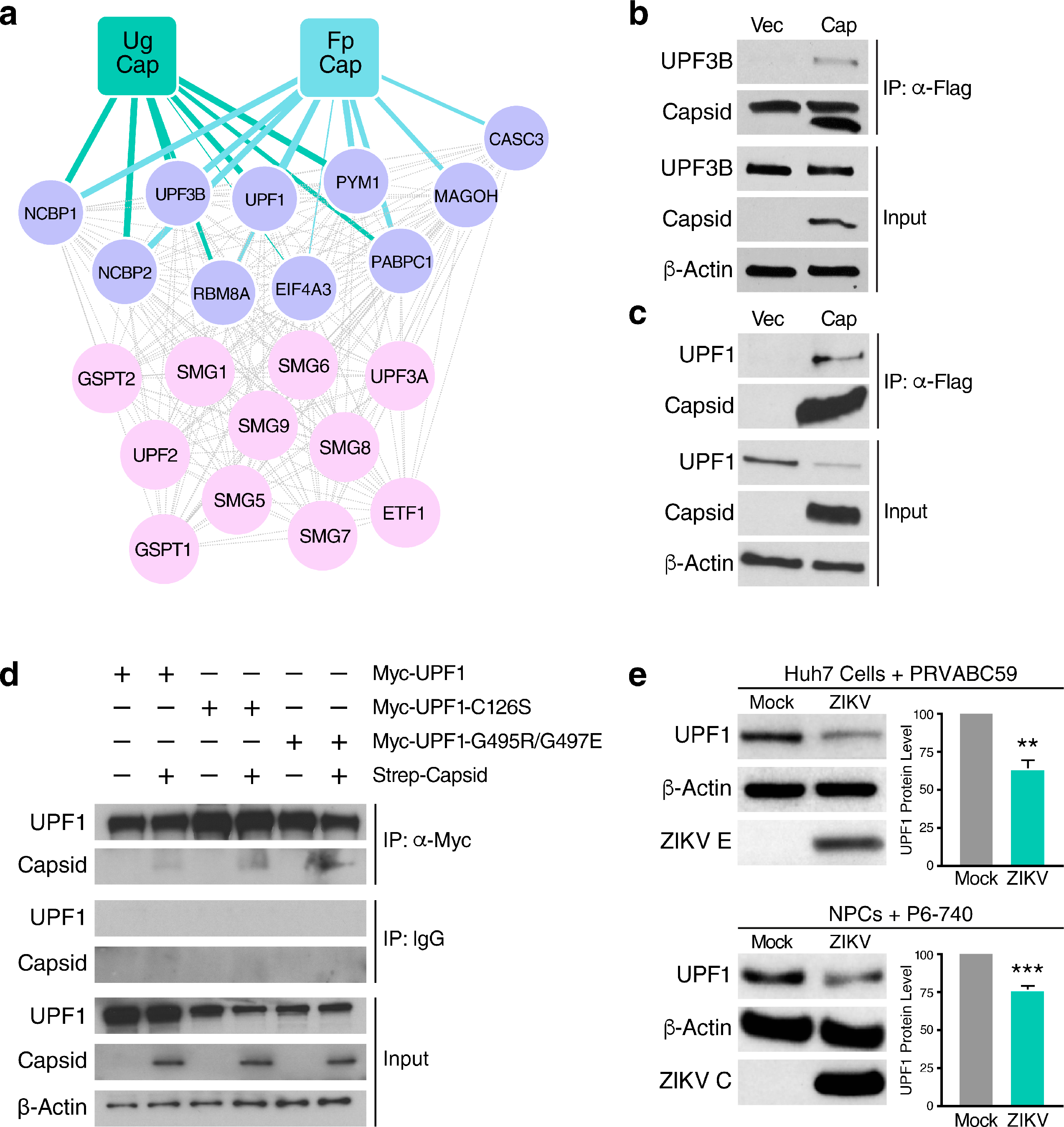
The capsid protein of ZIKV interacts with the NMD pathway. (a) Ugandan ZIKV capsid (Ug Cap, MR 766) and French Polynesian ZIKV capsid (Fp Cap, H/PF/2013) PPI maps that show significant enrichment for host NMD-associated factors (purple), as identified by AP-MS (SAINTq probability score > 0.9 and FDR < 0.05). Ten interactions between Fp Cap and host NMD factors (hypergeometrical test, *P* value = 7.16 x 10^−10^) and eight interactions between Ug Cap and host NMD factors (*P* value = 3.45 x 10^−7^) were identified. (b) Co-immunoprecipitation (co-IP) and western blot analysis of HEK293T cells transfected with vector or Flag-tagged ZIKV capsid (H/PF/2013, Asian lineage) and harvested at 48 hpt to immunoprecipitate endogenous UPF3B. The upper band detected in the IP Capsid blot represents a non-specific artifact. (c) Co-IP and western blot analysis of HEK293T cells transfected with vector or Flag-tagged ZIKV capsid and harvested at 48 hpt to immunoprecipitate endogenous UPF1. (d) Myc-tag co-IP and western blot analysis of HEK293T cells transfected with Strep-tagged ZIKV capsid and Myc-UPF1 (WT), Myc-UPF1-C126S (RNA-binding mutant), or Myc-UPF1-G495R/G497E (ATPase/helicase mutant) and harvested at 48 hpt to immunoprecipitate ZIKV capsid. (e) Western blot analysis of UPF1 levels in mock-infected and ZIKV-infected (PRVABC59, MOI of 1) Huh7 cells or mock-infected and ZIKV-infected (P6-740, MOI of 1) NPCs harvested at 48 hpi, with β-actin and ZIKV envelope (ZIKV E) or ZIKV capsid (ZIKV C) protein serving as loading and infection controls, respectively. Densitometric analyses were performed using ImageJ to quantify relative band intensities. Data are represented as mean ± s.e.m. *P* values were calculated by unpaired Student’s *t*-test. \*\**P* ≤ 0.01; \*\*\**P* ≤ 0.001. n= 3 independent experiments.

### ZIKV capsid degrades UPF1, the master regulator of NMD

Surprisingly, we consistently observed a decrease in UPF1, but not UPF3B, protein levels in the input lysate of ZIKV capsid-transfected cells, pointing to a specific perturbation of endogenous UPF1 expression by ZIKV capsid (Fig. 2c). To confirm that UPF1 protein levels are dysregulated during ZIKV infection, we performed western blot analysis of infected Huh7 cells and NPCs. Cellular UPF1 protein levels were consistently downregulated by ∼50% in ZIKV-infected Huh7 cells, whereas a ∼25% reduction was observed in ZIKV-infected NPCs (Fig. 2e), mirroring the difference in infection efficiencies achieved in these two cell systems. UPF1 transcript levels were not decreased in ZIKV-infected cells or following ZIKV capsid overexpression, indicating that UPF1 is post-transcriptionally downregulated during ZIKV infection (Supplemental Fig. 1a,b, respectively).

Because ZIKV capsid and UPF1 both localize to the nucleus and the cytoplasm (13, 40) we performed fractionation studies in ZIKV capsid-transfected HEK293T cells to determine if UPF1 is downregulated within a specific cellular compartment. Capsid expression markedly decreased nuclear UPF1 levels, whereas cytoplasmic levels were unchanged (Fig. 3a). We next examined a potential role for the autophagic and proteasomal pathways, both of which are known to mediate nuclear protein degradation, in ZIKV capsid-induced UPF1 downregulation. As shown in Supplemental Figure 2, nuclear UPF1 levels in ZIKV capsid-transfected cells were not rescued by inhibition of cellular autophagy via bafilomycin A1 treatment. However, nuclear UPF1 levels were restored in a dose-dependent manner following treatment with the proteasome inhibitor bortezomib (Fig. 3b), indicating enhanced proteasomal degradation of nuclear UPF1 in the presence of ZIKV capsid. Although ZIKV capsid colocalized with endogenous UPF1 in the cytoplasm of transfected Huh7-Lunet cells (Mander’s colocalization coefficient of ∼57%), we detected very little colocalization within the nucleus (∼7%) (Supplemental Figure 3). We hypothesized that this was due to specific degradation of UPF1 by ZIKV capsid in the nucleus. Indeed, when cells were treated with bortezemib, the fraction of nuclear UPF1 colocalizing with ZIKV capsid increased ∼8-fold, while the fraction of nuclear capsid interacting with UPF1 remained unchanged (Figure 3c). These results demonstrate that ZIKV capsid interacts with UPF1 both in the cytoplasm and nucleus, but specifically targets nuclear UPF1 for proteasomal degradation.

**Figure 3.**
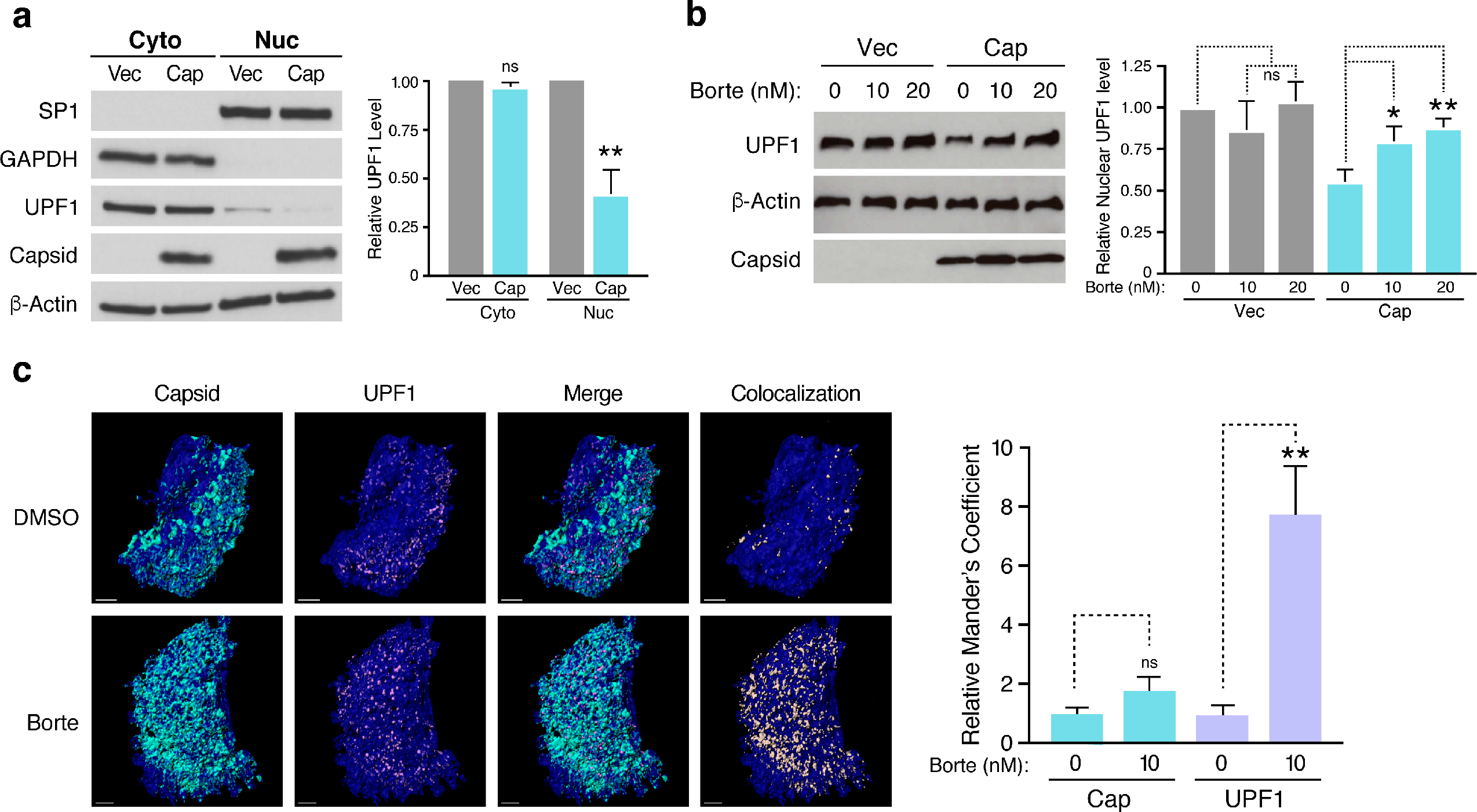
ZIKV capsid degrades UPF1, the master regulator of NMD, via a proteasome-dependent mechanism. (a) Western blot analysis of UPF1 levels in subcellular fractionated HEK293T cells transfected with vector or Flag-tagged ZIKV capsid (H/PF/2013, Asian lineage) for 48 h. GAPDH was used as a cytoplasmic marker and SP1 as a nuclear marker to ensure optimal fractionation. Densitometric analyses were performed using ImageJ to quantify relative band intensities. Data are represented as mean ± s.e.m. *P* values were calculated by unpaired Student’s *t*-test. \*\**P* ≤ 0.01; ns, not significant. n= 3 independent experiments. (b) Western blot analysis of nuclear UPF1 levels in fractionated HEK293T cells transfected with vector or Flag-tagged ZIKV capsid for 48 h. Cells were treated with DMSO or increasing concentrations of the proteasome inhibitor bortezomib for 24 h before harvest. Densitometric analyses were performed using ImageJ to quantify relative band intensities. Data are represented as mean ± s.e.m. *P* values were calculated by one-way ANOVA with multiple comparisons. \**P* ≤ 0.05; ns, not significant. n= 3 independent experiments. (c) Representative 3D confocal microscopy images of the nuclei of Huh7-Lunet cells transfected with Strep-tagged ZIKV capsid. Cells were treated at 24 hpt with DMSO or 10 nM bortezomib and processed for immunostaining at 48 hpt with antibodies against Strep-tag (turquoise) and endogenous UPF1 (purple). DAPI (blue) was used to stain and define the nuclei. Each channel was reconstructed digitally for visualization of the 3D colocalization. The thresholded Mander’s correlation coefficients were determined and *P* value was calculated by unpaired Student’s *t*-test. \*\**P* ≤ 0.01. n = 8 cells per condition. Scale bar represents 3 μm.

### UPF1 is a restriction factor of ZIKV

To test the effect of lowered UPF1 levels on ZIKV infection, we further decreased UPF1 expression prior to ZIKV infection by transfecting NPCs with either non-targeting siRNA or a pool of UPF1-specific siRNAs. We then infected the transfected cells with ZIKV and measured viral RNA levels, as well as infectious titers, 48 h post-infection (hpi). UPF1 knockdown was successful in siRNA-treated cells, as confirmed by western blot analysis (Fig. 4a). The depletion of UPF1 in NPCs prior to infection resulted in a significant increase in both ZIKV RNA levels and infectious virus production (Fig. 4b,c, respectively), indicating that expression of UPF1 restricts ZIKV infection at or before the RNA replication stage. To differentiate between these two stages, we analyzed double-stranded RNA (dsRNA) intermediates representing presumed viral RNA replication centers in infected NPCs (Fig. 4d)(41). Using confocal microscopy and 3D reconstruction analyses, we observed no significant difference in the number and size of dsRNA foci per cell when comparing ZIKV-infected, UPF1-depleted NPCs to ZIKV-infected cultures expressing UPF1 (Fig. 4e,f). Instead, we found a significant increase in the number of infected cells in NPC cultures when UPF1 was depleted, indicating that UPF1 regulates permissivity of NPCs to ZIKV infection at an early stage prior to viral RNA replication (Fig. 4g).

**Figure 4.**
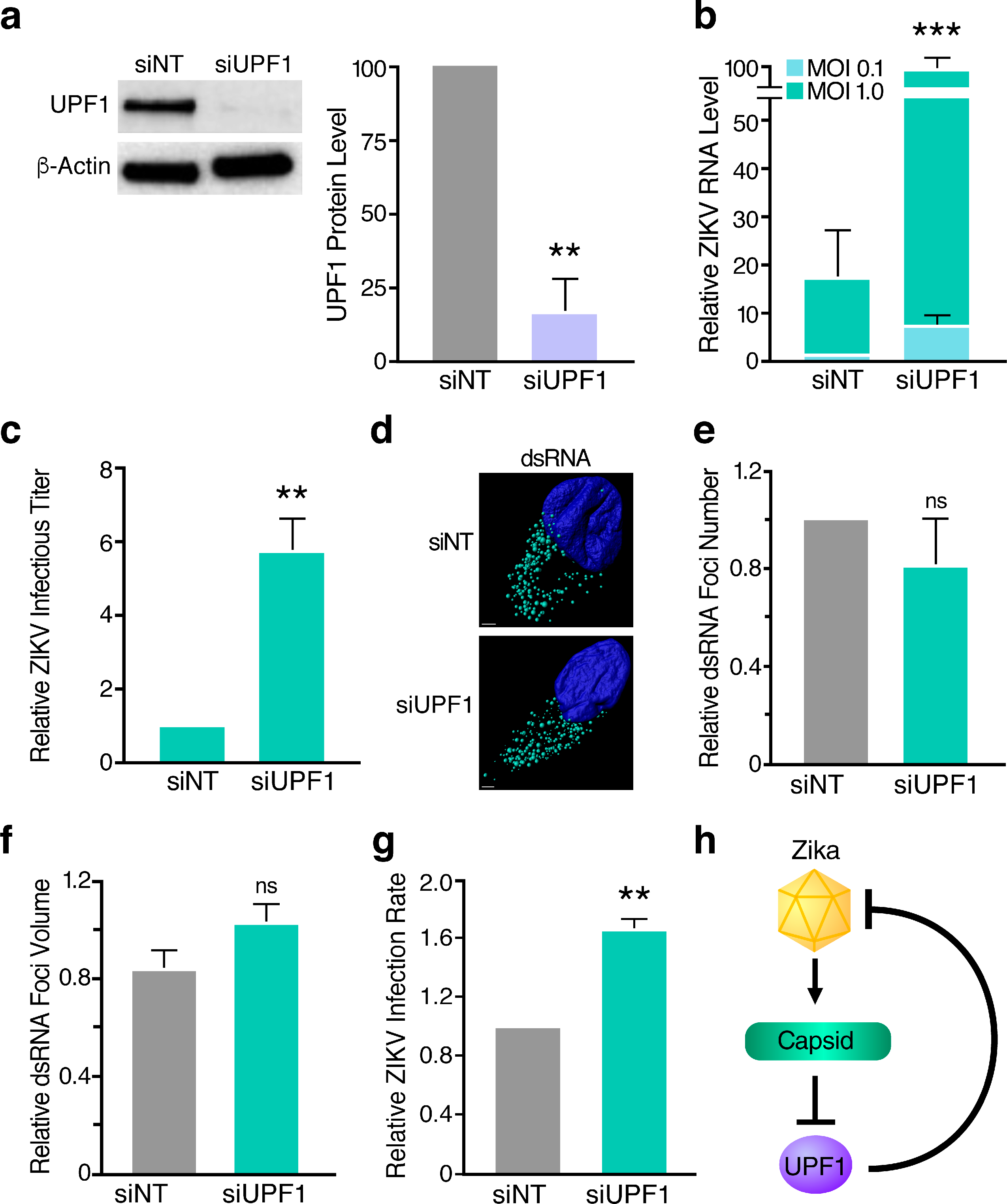
UPF1 knockdown enhances the permissivity of NPCs to ZIKV infection. (a) Western blot analysis of UPF1 levels in NPCs transfected with non-targeting siRNA (siNT) or a pool of UPF1-specific siRNAs (siUPF1) at 96 hpt. Densitometric analyses were performed using ImageJ to quantify relative band intensities. Data are represented as mean ± s.e.m. *P* value was calculated by unpaired Student’s *t*-test. \*\**P* ≤ 0.01. n= 3 independent experiments using one NPC line. (b) ZIKV RNA levels in siNT-treated or siUPF1-treated NPCs infected with ZIKV strain PRVABC59 at an MOI of 0.1 or 1 and harvested at 48 hpi. Data are represented as mean ± s.e.m. *P* value was calculated by two-tailed ratio paired Student’s *t*-test. \*\*\**P* ≤ 0.001. n= 3 independent experiments using one NPC line. (c) Released infectious virus from siNT-treated or siUPF1-treated, ZIKV-infected (MOI of 1) NPCs harvested at 48 hpi. Data are represented as mean ± s.e.m. *P* value was calculated by unpaired Student’s *t*-test. \*\**P* ≤ 0.01. n= 3 independent experiments using one NPC line. (d) Representative confocal microscopy images of a ZIKV-infected, siNT-treated NPC or a ZIKV-infected, siUPF1-treated NPC with the nuclei stained with DAPI (blue) and ZIKV dsRNA foci stained with the anti-dsRNA mAb J2 (teal). 3D image rendering and reconstructed dsRNA foci were produced using the Imaris spot detection function. Scale bar represents 2 μm. (e) Number of dsRNA foci were averaged for each cell. Data are represented as mean ± s.e.m. *P* value was calculated by two-tailed ratio paired Student’s *t*-test. ns, not significant. n = 3 independent experiments using two NPC lines, with 3-10 cells analyzed per condition for each experiment. (f) Measurement of dsRNA foci volume were averaged for each cell. Data are represented as mean ± s.e.m. *P* value was calculated by two-tailed ratio paired Student’s *t*-test. ns, not significant. n = 3 independent experiments using two NPC lines, with 3-10 cells analyzed per condition for each experiment. (g) Infection rates of siNT-treated or siUPF1-treated, ZIKV-infected (MOI of 1) NPCs measured at 48 hpi. Fixed cells were subjected to the anti-DENV mAb 1.6D, which also recognizes the ZIKV envelope protein (56). Data are represented as mean ± s.e.m. *P* value was calculated by two-tailed ratio paired Student’s *t*-test. \*\**P* ≤ 0.01. n = 3 independent experiments using two NPC lines. (h) Model of the interaction between the capsid protein of ZIKV and UPF1 of the NMD pathway.

## Discussion

In summary, we identified the NMD pathway as a restriction mechanism for ZIKV infection in human NPCs. NMD was partially inactivated in ZIKV-infected NPCs through expression of the viral capsid protein and the resulting degradation of host nuclear UPF1. As further weakening NMD by depleting UPF1 resulted in a marked increase in the number of infected cells, we propose a model in which an evolutionary “arms race” between cellular NMD and ZIKV determines whether a cell is successfully infected (Fig.4h).

Downregulation of UPF1 by ZIKV capsid is not complete, and is likely limited by the damaging effects of NMD disruption, as illustrated by the upregulation of genes involved in cell growth arrest and apoptosis. Indeed, knockout of UPF1 and other members of the NMD pathway is embryonic lethal in mice (19). However, mice haploinsufficient for NMD factors upstream of UPF1, including Magoh, Rbm8a, and Eif4a3, develop microcephaly (20–22). Thus, the reduction in nuclear UPF1 we observe in ZIKV-infected NPCs could contribute to the microcephaly phenotype caused by ZIKV infection in the fetal brain. While fetal and adult NPCs appear to be transcriptionally distinct (42), it has been shown that adult NPCs are also permissive to ZIKV infection (43). As the NMD pathway is a ubiquitous cellular surveillance mechanism, it is likely that ZIKV capsid targets UPF1 for degradation in any cell type that is susceptible to ZIKV infection. Accordingly, we have found that UPF1 is degraded following infection of both NPCs and hepatic Huh7 cells.

Why ZIKV capsid specifically downregulates nuclear UPF1, and how nuclear UPF1 contributes to ZIKV restriction remain unanswered. Several studies suggest that NMD is associated with the nucleus, although this issue remains controversial. Multiple transcripts, such as those encoding T cell receptor beta, triosephosphate isomerase and mouse major urinary protein, have been shown to be specifically degraded in purified nuclei or reduced in nuclear fractions (44). These data support the model that selectively depleting nuclear UPF1 levels disrupts NMD function in ZIKV-infected cells. In addition, UPF1 is involved in several other processes within the nucleus, including nuclear-associated RNA metabolism, cell cycle progression, and DNA replication (40). Therefore, by targeting nuclear UPF1, ZIKV could disrupt these processes and program target cells for viral replication. Notably, viral RNA replication is thought to occur solely within the cytoplasmic compartment (45, 46). Using confocal microscopy and 3D reconstruction, we did not detect dsRNA foci localized within the nuclei of ZIKV-infected cells (data not shown), supporting our finding that UPF1 does not restrict viral RNA replication. While our results suggest a role for nuclear UPF1 in ZIKV restriction, it is possible that UPF1 also serves as a restriction factor of ZIKV within the cytoplasm. Previously, it was shown that UPF1 suppresses alphavirus replication by degrading the incoming viral RNA following uncoating in the cytosol (28). Thus, ZIKV may possess an additional mechanism to prevent cytoplasmic UPF1 from targeting its incoming RNA genome for destruction.

Our data reveal that nuclear UPF1 is degraded by ZIKV capsid in a proteasome-dependent manner. While the nuclear proteasome has not been specifically linked to microcephaly, it plays critical roles in the regulation of chromatin structure, gene expression, DNA repair, and protein quality control (47). Thus, the co-opting of the nuclear proteasome by ZIKV capsid to degrade UPF1 could disrupt its normal proteasomal activity and further contribute to the cytopathic effects associated with ZIKV infection. Furthermore, given that the capsid protein of the closely related dengue virus can translocate across cell membranes (48), it is possible that capsid released from apoptotic, ZIKV-infected cells can enter neighboring, uninfected cells to degrade UPF1, thus increasing permissivity of bystander cells to ZIKV infection. Studies are ongoing to determine the precise molecular mechanism of ZIKV capsid-mediated UPF1 degradation and how UPF1 depletion enhances ZIKV replication, directly or indirectly. Ultimately, these data may help inform new therapeutic approaches, as reinforcement of the antiviral properties of the NMD pathway is expected to enhance resistance of NPCs to ZIKV infection and to promote normal neurodevelopment in infected fetuses.

## Materials and Methods

### Viruses and cells

Two Asian lineage strains of ZIKV, P6-740 (ATCC VR-1845) and PRVABC59 (ATCC VR-1843), were used for all experiments. ZIKV stocks were propagated in Vero cells (ATCC) and titers were determined by plaque assays on Vero cells. Huh7 cells (ATCC), Huh7-Lunet cells (Ralf Bartenschlager, Heidelberg University) and Vero cells were maintained in Dulbecco’s Modified Eagle’s Medium (DMEM) with 10% fetal bovine serum (FBS), 2 mM L-glutamine, 100 U/mL penicillin, and 100 μg/mL streptomycin. HEK293T cells (ATCC) were maintained in DMEM/H21 medium supplemented with 10% FBS, 100 U/mL penicillin, 100 μg/mL streptomycin, and 1 mM sodium pyruvate or DMEM with 10% FBS, 2 mM L-glutamine, 100 U/mL penicillin, and 100 μg/mL streptomycin. Human iPSC-derived NPCs were generated and maintained as described previously (49). All of the human fibroblast cell lines used to generate iPSCs came from the Coriell Institute for Medical Research and Yale Stem Cell Center. The iPSCs used in these studies were the CTRL2493nXX, CS2518nXX, and Cs71iCTR-20nXX lines. CTRL2493nXX was derived from the parental fibroblast line ND31845 that was biopsied from a healthy female at 71 years of age. CS2518nXX was derived from the parental fibroblast line ND30625 that was biopsied from a healthy male at 76 years of age. CS71iCTR-20nXX was derived from the parental fibroblast line ND29971 that was biopsied from a female at 61 years of age. For virus infections, NPCs plated on Matrigel-coated (Corning) multi-well plates or Huh7 cells were infected with ZIKV at a multiplicity of infection (MOI) of 0.1 or 1 for 2 h at 37°C. Infected cells were harvested at 48 hpi for all analyses.

### Affinity purification, mass spectrometry, and AP-MS scoring

The ZIKV capsid open reading frames (ORFs) from the Ugandan 1947 strain MR 766 or the French Polynesian 2013 strain H/PF/2013 were cloned into pCDNA4_TO with a C-terminal 2xStrep II affinity tag for expression in human cells. The viral capsid proteins (three biological replicates), as well as GFP (two biological replicates) and empty vector (ten biological replicates) as negative controls, were expressed in HEK293T cells and affinity purifications were performed as previously described (50). Briefly, clarified lysates were incubated with Strep-Tactin Superflow (IBA) overnight at 4HC. Proteins were eluted with 50 mM Tris pH 7.5, 150 mM NaCl, 1 mM EDTA containing 2.5 mM Desthiobiotin (IBA) for 30 min at 44C. Lysates and affinity purified eluates were analyzed by western blot and silver stain PAGE to confirm expression and purification. Purified protein eluates were digested with trypsin for LC-MS/MS analysis. Samples were denatured and reduced in 2M urea, 10 mM NH_4_HCO_3_, 2 mM DTT for 30 min at 60°C, then alkylated with 2 mM iodoacetamide for 45 min at room temperature. Trypsin (Promega) was added at a 1:100 enzyme:substrate ratio and digested overnight at 37°C. Following digestion, samples were concentrated using C18 ZipTips (Millipore) according to the manufacturer’s specifications. Peptides were resuspended in 15 μL of 4% formic acid and 3% ACN, and 1-2 μL of sample was loaded onto a 75 μm ID column packed with 25 cm of Reprosil C18 1.9 μm, 120Å particles (Dr. Maisch GmbH). Peptides were eluted into a Q-Exactive Plus (Thermo Fisher) mass spectrometer by gradient elution delivered by an Easy1200 nLC system (Thermo Fisher). The gradient was from 4.5% to 32% acetonitrile over 53 min. All MS spectra were collected with oribitrap detection, while the 20 most abundant ions were fragmented by higher energy collisional dissociation (HCD) and detected in the orbitrap. All data was searched against the SwissProt Human protein sequences, combined with ZIKV sequences and GFP. Peptide and protein identification searches, as well as label-free quantitation, were performed using the MaxQuant data analysis algorithm and all peptide and protein identifications were filtered to a 1% false-discovery rate (51, 52). SAINTq (53) was used to calculate the probability of bait-prey interactions for both Ugandan ZIKV capsid and French Polynesian ZIKV capsid against the negative controls, including GFP and empty vector, with protein intensities as input values. We applied a combined threshold of probability of interaction (AvgP) greater than 0.90 and Bayesian False Discovery Rate of less than 0.05.

### Quantitative real-time reverse transcription-PCR (qRT-PCR)

Total cellular RNA was isolated from Huh7 cells and NPCs using the RNeasy Mini Kit (Qiagen). cDNA was synthesized with oligo(dT)_18_ (ThermoFisher Scientific) primers, random hexamer (Life Technologies) primers, and AMV reverse transcriptase (Promega). The cDNA was then used in SYBR Green PCR Master Mix (ThermoFisher Scientific) according to manufacturer’s instructions and analyzed by qPCR (Bio-Rad ABI 7900). The primers used for ASNS, CARS, SC35 1.7, GAPDH, HPRT1, LDHA, and 18S rRNA have been described previously (29). The additional primers used were ZIKV PRVABC59 forward primer 5’-GAG ACG AGA TGC GGT ACA GG −3’, ZIKV PRVABC59 reverse primer 5’-CGA CCG TCA GTT GAA CTC CA −3’, UPF1 forward primer 5’-CTG CAA CGG ACG TGG AAA TAC −3’, UPF1 reverse primer 5’-ACA GCC GCA GTT GTA GCA C −3’, DDIT3 forward primer 5’-CTG CTT CTC TGG CTT GGC TG −3’, DDIT3 reverse primer 5’-GCT CTG GGA GGT GCT TGT GA −3’, GADD45A forward primer 5’-GAG CTC CTG CTC TTG GAG AC −3’, GADD45A reverse primer 5’-GCA GGA TCC TTC CAT TGA GA −3’, GADD45B forward primer 5’-TGA CAA CGA CAT CAA CAT C −3’, and GADD45B reverse primer 5’-GTG ACC AGA GAC AAT GCA G −3’. Relative levels of each transcript were normalized by the delta threshold cycle method to the abundance of 18S rRNA or GAPDH, with mock-infected cells or vector-transfected cells set to 1.

### Western blot analysis

Cells were lysed in RIPA lysis buffer (50 mM Tris-HCl, pH 8, 150 mM NaCl, 1% NP-40, 0.5% sodium deoxycholate, 0.1% SDS, supplemented with Halt™ protease inhibitor cocktail (ThermoFisher Scientific)) to obtain whole cell lysates or lysed using the NE-PER nuclear and cytoplasmic extraction kit (ThermoFisher Scientific) to obtain cytoplasmic and nuclear fractions. Proteins were separated by SDS-PAGE and transferred to nitrocellulose membranes (Bio-Rad). Blots were incubated with the indicated primary antibody: anti-UPF3B (ab134566, Abcam), anti-UPF1 (12040, Cell Signaling Technology, Inc.), anti-ZIKV Capsid (C) (GTX133304, GeneTex), anti-Flag (F7425, Sigma-Aldrich), anti-β-actin (A5316, Sigma-Aldrich), anti-ZIKV Envelope (E) (GTX133314, GeneTex), anti-SP1 (sc-14027, Santa Cruz Biotechnology), anti-GAPDH (5174, Cell Signaling Technology, Inc.), anti-Myc tag (ab9106, Abcam), anti-Strep tag (ab18422, Abcam), and anti-p62 (ab56416, Abcam). Proteins were visualized by chemiluminescent detection with ECL and ECL Hyperfilm (Amersham). Differences in band intensity were quantified by densitometry using ImageJ.

### Immunoprecipitations

Cells were lysed in either RIPA lysis buffer or IP lysis buffer (150mM NaCl, 50mM Tris pH 7.4, 1mM EDTA, 0.5% NP-40 substitute, supplemented with Halt™ protease inhibitor cocktail (ThermoFisher Scientific) at 4°C and passed through a G23 needle. Clarified lysates were immunoprecipitated with Flag M2 agarose (Sigma), anti-Myc tag (ab9106, Abcam), or normal rabbit IgG (sc-2027, Santa Cruz Biotechnology) overnight, washed in lysis buffer, and resuspended in Laemmli buffer for SDS-PAGE. Western blot analysis of immunoprecipitated proteins was performed as described above.

### Immunofluorescence

Transfected Huh7-Lunet cells or infected NPCs were collected at 48 h and plated onto 22 × 22 mm #1.5 coverslips. Cells were fixed in 4% paraformaldehyde, permeabilized with 0.1% Triton X-100, and blocked in 3% bovine serum albumin. Cells were then immunostained with the indicated antibodies: anti-Strep Tag (Abcam, ab184224), anti-UPF1 (Abcam, ab109363), human anti-DENV mAb 1.6D (Sharon Isern and Scott Michael, Florida Gulf Coast University), which recognizes the ZIKV envelope protein, anti-dsRNA mAb J2 (SCICONS), and the appropriate fluorophore-conjugated secondary antibodies. Coverslips were mounted onto glass slides using Vectashield^®^ Mounting Medium with DAPI (Vector Laboratories) and analyzed by fluorescence microscopy (Zeiss Axio Observer ZI) or confocal microscopy (Zeiss LSM 880). For acquiring high-resolution images, cells were imaged on the Zeiss LSM 880 with Airyscan using a 20x/0.8 or 63x/1.4 M27 oil immersion objective. A total of 15-20 (20x objective) or 60-80 (63x objective) Z-slices were acquired every 0.88 μm or 0.15 μm, respectively. The resulting Z-stack was reconstructed and rendered in 3D using Imaris software (Bitplane). Viral dsRNA foci were reconstructed via the Imaris spot detection function, which provided an analysis of total number and mean volume of foci within a cell, for images acquired using the 20x objective. Strep-tagged ZIKV capsid, UPF1, and dsRNA channels acquired using the 63x objective were reconstructed using the Imaris surfaces package. The Imaris colocalization function was used to determine overlap of fluorescence. Thresholding for background fluorescence was determined by the Imaris automatic thresholding tool that utilizes the Costes approach (54). The thresholded Mander’s correlation coefficient (MCC) measures the fraction of voxels with fluorescence positive for one channel that also contain fluorescence from another channel. The MCC is typically more appropriate for analysis of three-dimensional colocalization (55).

### Statistical analysis

Statistical differences between groups were analyzed using either a two-tailed unpaired Student’s *t*-test or a two-tailed ratio paired Student’s *t*-test, as stated in the figure legends. Hypergeometrical tests were used to calculate the probability of an overlap in gene dysregulation between ZIKV-infected NPCs and UPF1-depleted cells and to calculate the probability of ZIKV capsid bait-prey interactions. Data are represented as mean ± s.e.m. Statistical significance was defined as \**P* ≤ 0.05, \*\**P* ≤ 0.01, \*\*\**P* ≤ 0.001, and \*\*\*\**P* ≤ 0.0001.

## Acknowledgements

The authors would like to thank all members of the Ott laboratory, as well as Roman Camarda, Marius Walter, and Anna Maurer for helpful discussions and advice throughout the preparation of this manuscript. We thank Chia-Lin Tsou, the Gladstone Stem Cell Core, and Meredith Calvert from the Gladstone Microscopy Core for technical assistance, and Ralf Bartenschlager (Heidelberg University), Lynne Maquat (University of Rochester), and Sharon Isern and Scott Michael (Florida Gulf Coast University) for reagents. We are grateful to Veronica Fonseca for administrative support, John Carroll and Teresa Roberts for graphical design, and to Kathryn Claiborn, Eric Martens, and Gary Howard for editorial assistance. This work was supported by NIH/NIAID F32AI112262 to P.S.S., NIH/NINDS R01 NS101996-01 to S.F., NIH/NIAID U19AI1186101 to N.J.K., DOD/DARPA HR0011-11-C-0094 (PROPHECY) to N.J.K., NIH/NIAID R01 AI097552 to M.O., and the James B. Pendleton Charitable Trust.

---

**Supplemental Figure 1.**
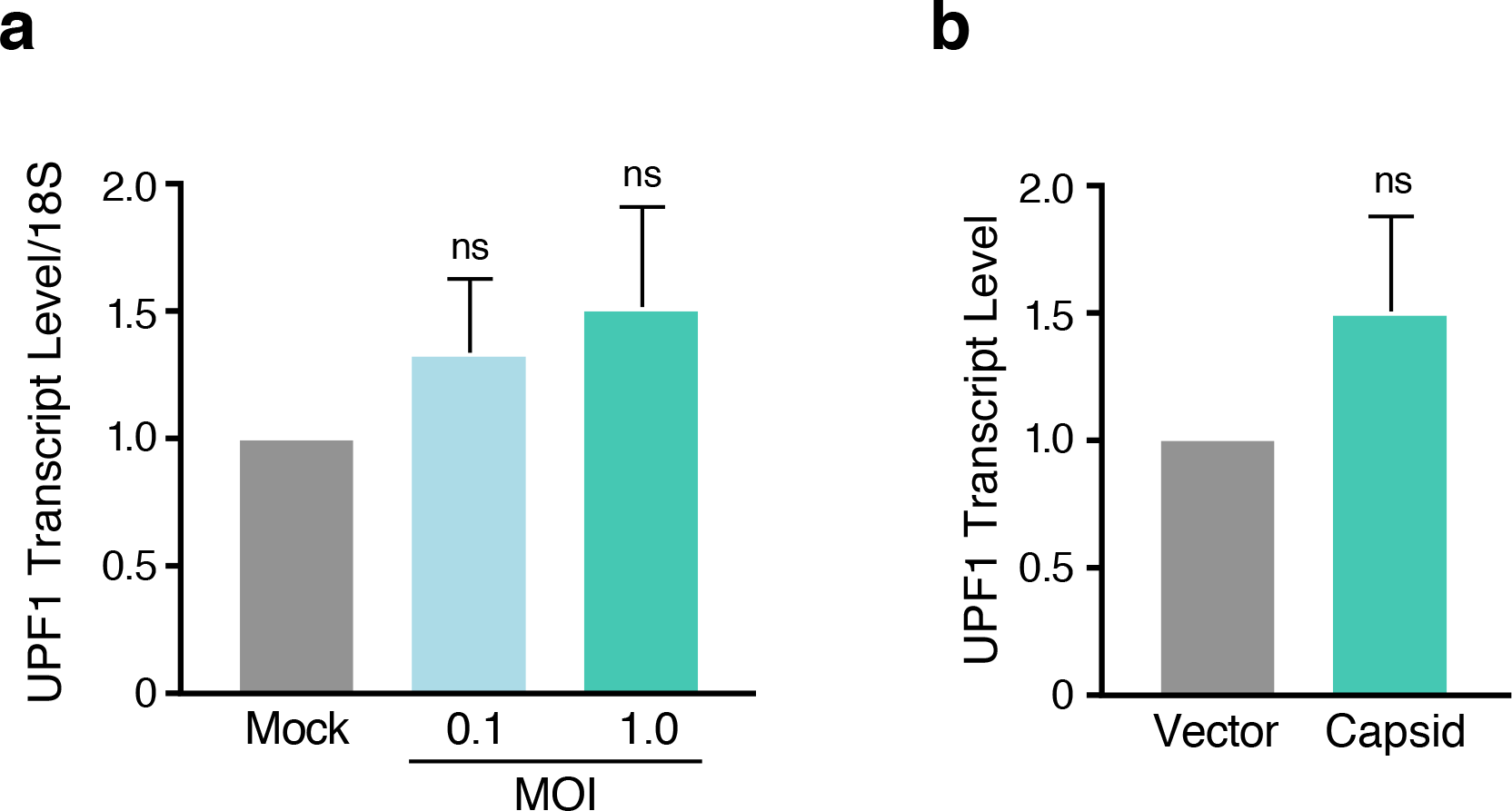
UPF1 is not transcriptionally downregulated during ZIKV infection or following ZIKV capsid overexpression. (a) UPF1 transcript levels from Huh7 cells mock-infected or infected with ZIKV strain PRVABC59 at an MOI of 0.1 or 1 and harvested at 48 hpi. Data are represented as mean ± s.e.m. *P* values were calculated by unpaired Student’s *t*-test. ns, not significant. n= 3 independent experiments. (b) UPF1 transcript levels from HEK293T cells transfected with vector or Strep-tagged ZIKV capsid (H/PF/2013, Asian lineage) and harvested at 48 hpt. Data are represented as mean ± s.e.m. *P* values were calculated by unpaired Student’s *t*-test. ns, not significant. n= 3 independent experiments.

**Supplemental Figure 2.**
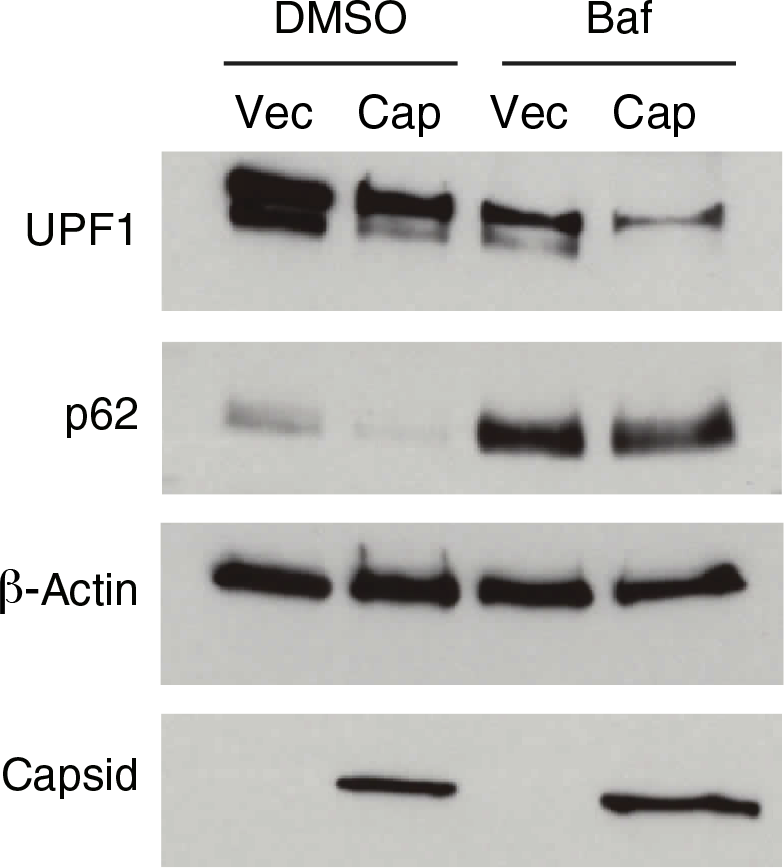
ZIKV capsid-induced UPF1 degradation is not dependent on autophagy. Western blot analysis of nuclear UPF1 levels in fractionated HEK293T cells transfected with vector or Flag-tagged ZIKV capsid for 48 h. Cells were treated with DMSO or the autophagy inhibitor bafilomycin A1 (Baf) (10 nM) for 24 h before harvest. Levels of p62, which is degraded by autophagy, were monitored to confirm autophagic inhibition following bafilomycin A1 treatment.

**Supplemental Figure 3.**
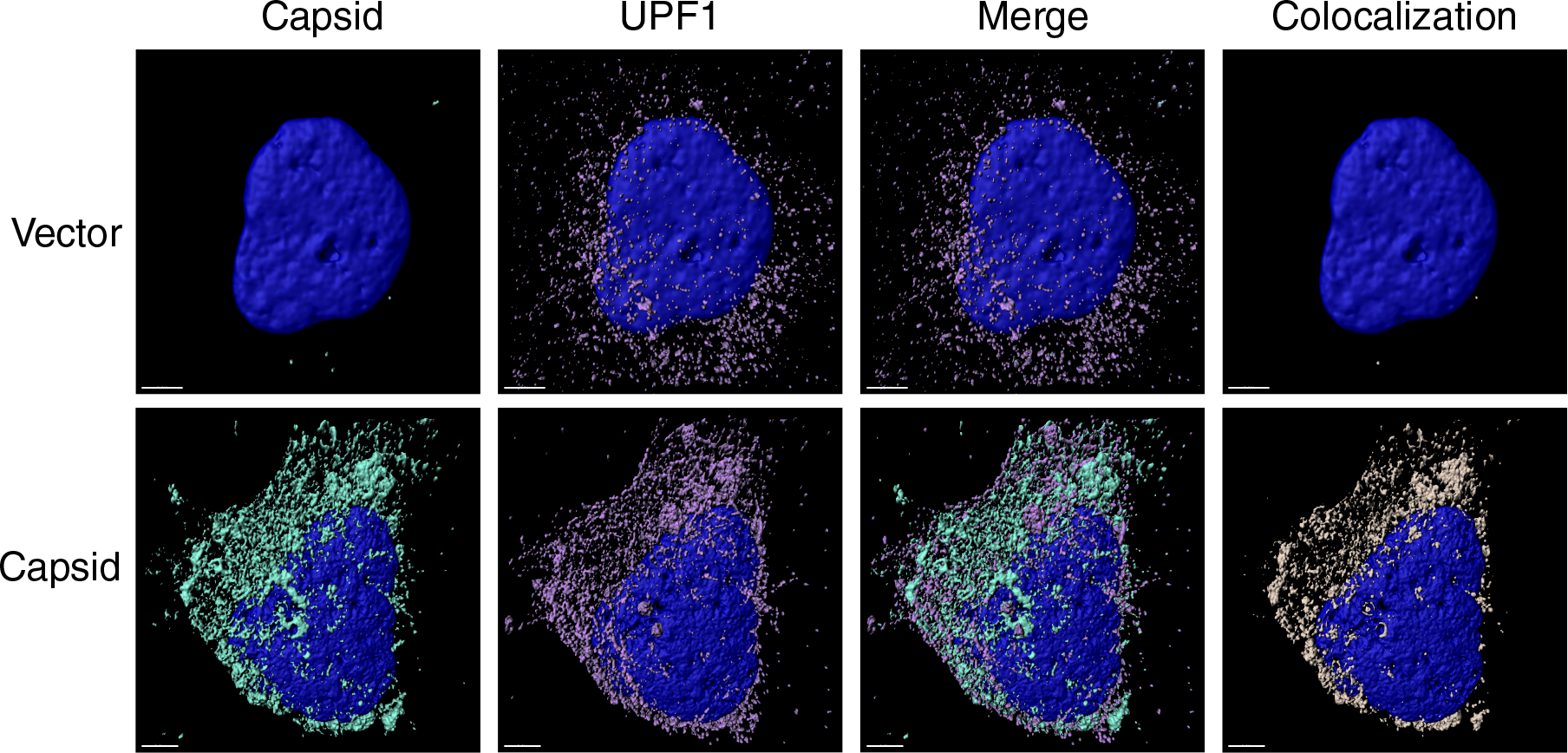
ZIKV capsid colocalizes with endogenous UPF1. Representative 3D confocal microscopy images of Huh7-Lunet cells transfected with vector or Strep-tagged ZIKV capsid. Cells were processed for immunostaining at 48 hpt and probed with antibodies against Strep-tag (turquoise) and endogenous UPF1 (purple). DAPI (blue) was used to stain the nuclei. Each channel was reconstructed digitally for visualization of the 3D colocalization. The thresholded Mander’s correlation coefficient for ZIKV capsid was 0.57 (n = 17), indicating that approximately 57% of the voxels positive for capsid fluorescence were also positive for UPF1 fluorescence. Scale bar represents 5 μm.

